# Summed contrasts can conceal reversed regional preferences: directional and dose criteria for screening transcranial direct current stimulation montage pairs

**DOI:** 10.64898/2026.10.07.757440

**Authors:** Dylan Bruschi, Ivanna Joseph, Adam Schiavi

**Author notes:** Corresponding author. E-mail address (D. Bruschi). Affiliations identify the institutions at which the work was carried out, not necessarily current appointments. I.J. completed Johns Hopkins neurocritical-care fellowship training in 2026; A.S. holds a faculty appointment in the School of Medicine.

## Abstract

**Background:** The regional-matching design evaluated here requires two transcranial direct current stimulation (tDCS) montages, each stronger in its own target region, with adequate exposure. Summed contrasts can be positive when only one preference holds.

**Objective:** To test whether a summed double-dissociation index (DDI), overlap and cohort-median dose rules identify such pairs, for a focused-attention versus open-monitoring experiment.

**Methods:** Twenty-four montage pairs were simulated in 20 anatomical head models (SimNIBS 4.6, 2 mA). A four-rule screen (DDI, overlap, cohort-median dose) was recorded before cohort results; a post hoc conjunction required both regional contrasts positive in ≥80% of heads. Two leading pairs were compared on per-head dose, a second solver and 60 conductivity stress-test draws.

**Results:** Eighteen pairs passed the screen, but 13 failed the conjunction; one with positive median DDI (0.782) reversed the salience-region preference in all 20 heads. The nominated pair (F3-centered 4×1 ring; F8–Fp2 sponges; median DDI 1.561; both preferences in 20/20 heads) brought both anatomical targets to the 0.20 V/m operational floor in 11/20 heads; a conventional sponge pair (DDI 0.627) did so in 20/20, though both passed the cohort-median dose rule. The nominated pair’s separation held across a second solver and all 60 stress-test draws, but both network targets reached the floor in 19/60.

**Conclusions:** Summed indices and cohort-median dose rules can pass pairs that fail reciprocal-exposure criteria. Such screens should prespecify the directional conjunction and report per-head joint dose coverage. These are modeled exposures, not physiological or behavioral effects.

## 1. Introduction

For a regional-matching design that compares two transcranial direct current stimulation (tDCS) montages, the intended exposure reversal is explicit: montage A should expose region X more than montage B does, while B should expose region Y more than A does. Current-flow modeling is used to choose such pairs before participants are stimulated [1,2]. A screen may summarize each candidate pair with a single separation index, an overlap measure and a cohort-median dose criterion, as ours originally did. Each summary can hide a failure of the intended exposure manipulation: a summed contrast can be positive when one of the two required preferences is reversed, and a cohort-median dose can clear a floor while many individual heads fall below it.

Our application is a planned experiment pairing tDCS with two mindfulness practices. Focused attention (FA) trains sustained attention, detection of distraction and reorientation; open monitoring (OM) trains non-reactive monitoring of ongoing experience without a single object [3,4]. FA imaging findings implicate executive-control regions [5,6], whereas experiential self-reference and interoceptive-attention studies motivate a hypothesized insular emphasis for OM [7,8]. These are overlapping network emphases, not one-to-one assignments of practices to circuits. Prior work motivates pairing stimulation with contemplative practice [9,10,11], but tDCS effects depend on brain state, connectivity, anatomy and the complete field distribution, not on electrode labels alone [12,13,14].

The planned behavioral test predicts a reversal: montage A should outperform B during FA, and B should outperform A during OM. The regional-matching hypothesis being evaluated assumes a corresponding exposure reversal. The FA candidate must exceed the OM candidate in a control-region mask, and the OM candidate must exceed the FA candidate in a salience-region mask, in individual heads and above the operational dose floor. These are model-selection criteria for this hypothesis; other state-dependent or distributed mechanisms could produce behavioral interactions without this p99 pattern. A positive aggregate index or an electrode label does not establish the intended exposure manipulation.

We screened 24 candidate pairs in 20 anatomical head models with a four-rule screen recorded before the cohort results, then audited it against a directional conjunction and per-head dose coverage. We asked how many pairs that passed the summed screen failed the conjunction, whether separation and individual dose coverage favored the same pairs, and which conclusions survived a second solver and conductivity variation. The principal contribution is methodological: screening paired montages for reciprocal regional exposure, illustrated on a worked example. No participant was stimulated, and modeled exposure is not evidence of a physiological or behavioral effect.

## 2. Methods

### 2.1 Head models and simulations

We used 20 existing anatomical head models from Berger et al. [15]: de-identified, facially masked headreco models [16] of Human Connectome Project S1200 participants [17], distributed under CC BY 4.0 [18]. Ten female and ten male source records were drawn with a fixed seed, stratified by recorded sex and age band, after a quality-control exclusion; three further heads, disjoint from the evaluation set, served the calibration and sensitivity analyses (Supplementary Section S1). Primary finite-element solutions used SimNIBS 4.6 [1], solved at 1 mA and scaled linearly to 2 mA. Each montage was solved separately; a pair denotes a comparison between two separately solved montages, not simultaneous delivery. Fields were sampled on each head’s gray-matter voxel grid.

The analysis snapshot is tdcs-modeling commit 0209ed2 (tag v1.38-manuscript), deposited publicly [19]. Results from the same simulations also appear in a companion framework preprint [20]; they are not an independent replication, and this article is the dedicated methods-and-results report.

### 2.2 Candidate montages

Four FA and six OM candidates formed 24 pairs (Table 1). Conventional montages used 50 × 50 mm sponge footprints with a 40 × 40 mm inset rubber element. The nominated OM montage, OM-02, placed the anode at F8 and the return at the ipsilateral supraorbital site Fp2; the conventional alternative combined FA-02 (F3–Fp2) with OM-07 (FT8–FCz).

**Table 1.** The 10 candidates forming the 24 FA-OM pairs.

| **Candidate** | **Anode / target** | **Return** | **Geometry and status** |
|---|---|---|---|
| FA-01 | F3 | F4 | 50 × 50 mm sponges |
| FA-02 | F3 | Fp2 | 50 × 50 mm sponges |
| FA-05 | F3 | Four surrounds | 12 mm discs; nominal 35 mm ring |
| FA-06 | Control-mask array | Distributed | Locked optimization; defective off-target penalty |
| OM-01 | F8 | Fp1 | 50 × 50 mm sponges |
| OM-02 | F8 | Fp2 | 50 × 50 mm sponges |
| OM-04 | FT8 | Fp1 | 50 × 50 mm sponges |
| OM-05 | F8 | Four surrounds | 12 mm discs; nominal 35 mm ring |
| OM-06 | Salience-mask array | Distributed | Locked optimization; defective off-target penalty |
| OM-07 | FT8 | FCz | 50 × 50 mm sponges; reinstated post hoc |

Results are reported at 2 mA total source current. Ring surrounds share one return channel. FA/OM prefixes identify intended experimental roles, not demonstrated practice specificity.

The high-definition rings (FA-05 centered at F3; OM-05 at F8) used a 12 mm central disc and four 12 mm surrounds at a nominal 35 mm radius, implemented with the SimNIBS expand_to_center_surround procedure. The surrounds shared one return channel, so equal per-surround currents were not imposed, and the achieved electrode coordinates define the solved array.

FA-06 and OM-06 were locked optimized arrays intended for focality-constrained distributed targeting [21]. A configuration defect removed the off-target penalty, so they are reported as locked arrays, not focality-optimized solutions. Corrected arrays were analyzed separately as exploratory candidates (Supplementary Section S6).

### 2.3 Regional definitions

The recorded screen used lateralized frontal subsets of the Schaefer 400-parcel, seven-network atlas [22]: left control-network labels (C) and right salience/ventral-attention labels (S). These are frontal masks, not complete networks; label rules are given in Supplementary Section S1.

Anatomical dose used HCP-MMP1 parcels [23]: left middle-frontal cortex (46, a9-46v, p9-46v, 9-46d and 8Av) for FA and strict right anterior insula (AVI and AAIC) for OM. Dorsal anterior cingulate cortex (dACC; bilateral a24pr, p24pr, a32pr, 33pr and p32pr) was summarized separately as a salience-network node, not an off-network control [24,25].

After cohort results were available, cortical labels were capped to gray matter within 3.5 mm of the central surface, removing a median subcortical fraction of 23% from the strict-insula mask. This post hoc correction left electrode arrays, focality and Dice unchanged; its effect on the passing count is reported in Section 3.1.

### 2.4 Field summaries, directional contrasts and overlap

Here, regional dose denotes a modeled exposure summary: the volume-weighted within-region 99th percentile of field magnitude (p99). Reported doses are cohort medians unless stated otherwise. Field magnitude does not determine excitation, inhibition or engagement, and this percentile is not the fraction of a region engaged.

For head h, FA candidate a and OM candidate b, the two intended regional contrasts were

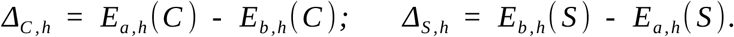

and the original double-dissociation index (DDI) normalized their sum:

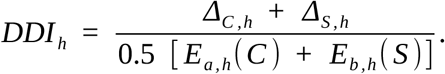

where E is the within-region p99. Directional refers to the sign of the between-montage magnitude contrast, not to electric-field vector orientation or neuronal polarization. A positive DDI can arise when one contrast is negative, because the denominator scales the sum without enforcing a directional conjunction.

Spatial overlap was the Dice coefficient between gray-matter maps above 50% of each montage’s robust peak, its 99.9th-percentile whole-gray-matter field; focality was the volume of each map. A Dice value of zero describes overlap at this relative threshold, not disjoint fields at all intensities.

### 2.5 Screening rules and audit measures

Four progression rules were recorded on 16 September 2026, before the cohort results: both cohort-median own-target network p99 values at least 0.20 V/m at 2 mA; positive DDI in at least 80% of heads; cohort-median Dice at most 0.50; and median DDI above 0.075311, a bar calibrated from simulated electrode-placement error (Section 2.6). The 0.20 V/m value is an operational development floor applied to an upper percentile, not an efficacy or neuronal threshold. Intracranial measurement/model comparisons [26,27] do not validate this percentile-based development rule. A cohort-median pass does not require every head to meet it.

The record did not precede all development information. Template results shaped the robust-peak and calibration definitions, OM-07 was reinstated after cohort results, and the cortical cap followed them; the amendment ledger is in Supplementary Section S1.

The audit used two further measures. A directional conjunction, added after the original results were inspected, required both ΔC > 0 and ΔS > 0 in at least 80% of heads; it was not used to redefine original passes. Per-head joint dose coverage counted heads in which both targets of a pair reached 0.20 V/m. Post hoc mean-field and anatomical node-mean variants tested whether conclusions depended on the p99 summary (Supplementary Section S2).

### 2.6 Sensitivity analyses

The DDI bar was the 90th percentile of absolute DDI change across 300 simulated placement perturbations of two anchor pairs in three held-out heads (5 mm Gaussian displacement, truncated at 15 mm; Supplementary Section S3). ROAST 4.0 [2] re-solved every montage except the optimized arrays, using its own 10–10 placement for sponges and the SimNIBS-solved centers for rings, so the ring comparison holds placement fixed (Supplementary Section S4). The archived conductivity stress test used 30 log-uniform Latin-hypercube settings in each of two held-out heads, giving 60 draws per pair. Its historical ranges were attributed to McCann, Pisano, and Beltrachini [28]; their correction excluded high-frequency studies [29]. We retained those archived results and separately summarized the existing draws within the corrected review’s tissue ranges (Supplementary Section S8). Central-surface sampling was compared with voxel sampling in two evaluation heads (Supplementary Section S5). None of these descriptive analyses determined the original gate.

### 2.7 Analysis and reproducibility

The unit of analysis was the anatomical head. We report cohort medians, exact numerators and denominators and complete candidate-pair results; we did not treat solver agreement as statistical independence, conductivity draws as participants or post hoc selection as confirmation. The directional conjunction used unrounded head-level contrasts.

Figures plot deposited head-level values at 2 mA and baseline conductivities; drawing them involved no new solve. The archived analysis code was developed with generative AI assistance under D.B.’s direction, as described in Supplementary Section S1. For this submission revision, ChatGPT (OpenAI; GPT-6 Astra Pro) assisted the numerical audit, the post hoc analyses in Supplementary Section S8, and the plotting code supplied with the submission. These calculations used recorded outputs and did not independently validate the finite-element solutions.

## 3. Results

### 3.1 The summed screen passed 18 pairs; 13 of them failed the directional conjunction

Eighteen of the 24 pairs passed the recorded screen (Table S1). The four pairs containing OM-05 failed the network-dose rule, and two pairs exceeded the overlap ceiling by margins (0.015 and 0.035) smaller than the 0.058 placement-related Dice change. The passing count was 14 before the post hoc cortical cap, which moved OM-06’s median network p99 to 0.201 V/m and added its four pairs; OM-06 nonetheless reached the anatomical anterior-insula floor in 0/20 heads.

Only five of the 18 passing pairs met the directional conjunction: FA-01 × OM-07, FA-02 × OM-04, FA-02 × OM-07, FA-05 × OM-02 and FA-05 × OM-06. The other 13 passed a screen built on a positive summed index without producing both intended preferences in at least 80% of heads.

### 3.2 A positive summed index concealed a reversed regional preference

FA-02 × OM-02 shows how the summed index failed. It passed with median DDI 0.782 and Dice 0.203. Its median control-region contrast was +0.290 V/m, but its median salience-region contrast was −0.063 V/m, and FA-02 produced the larger salience-mask p99 in every head (Fig. 1).

**Fig. 1.**
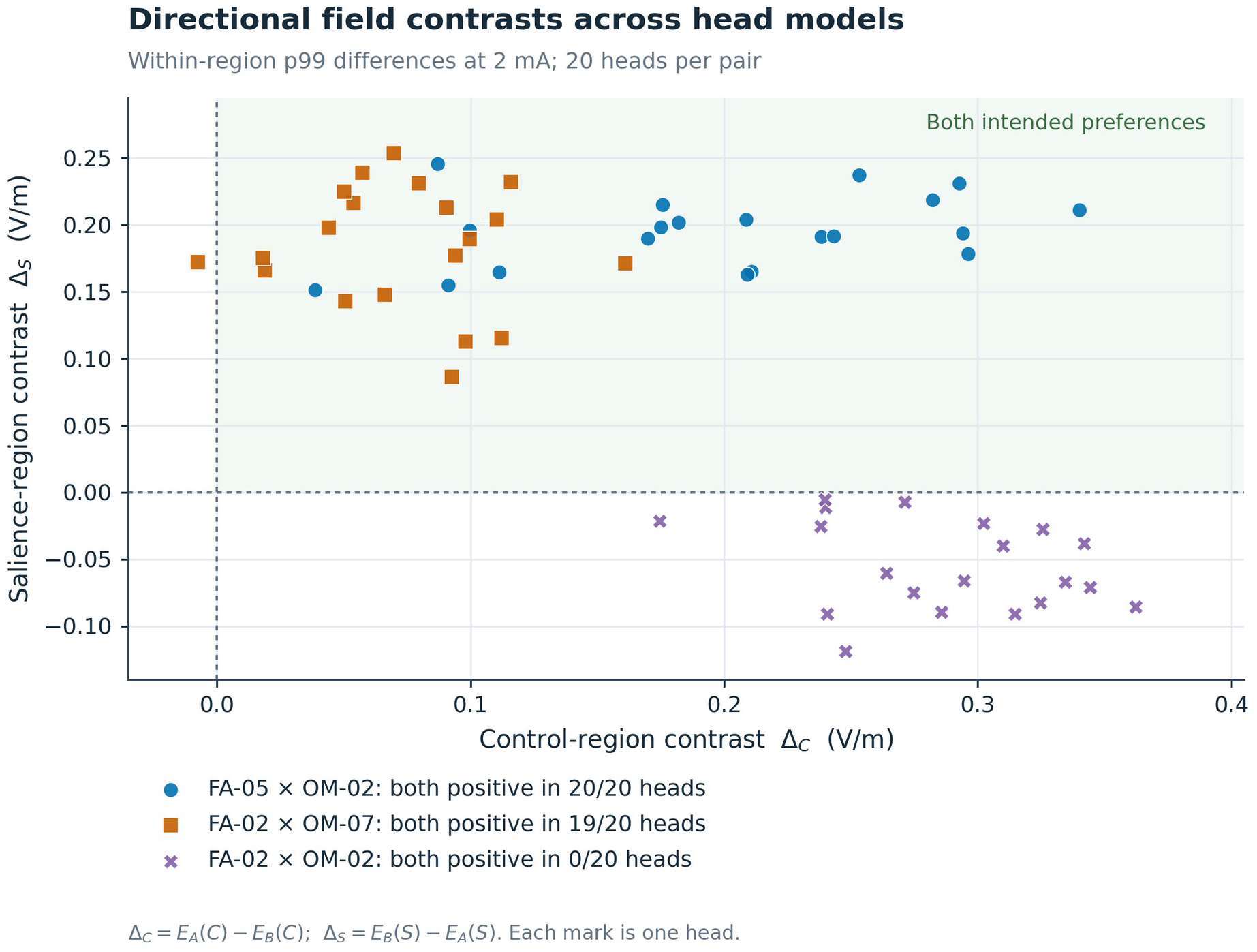
Directional contrasts by head. Each point is one head under one montage pair (20 heads per pair); the upper-right quadrant meets both intended preferences. FA-05 × OM-02 does so in 20/20 heads and FA-02 × OM-07 in 19/20. FA-02 × OM-02 has a positive median DDI but a negative salience-region contrast in every head. Values are differences in within-region p99 at 2 mA and baseline conductivities; no confidence intervals are shown.

The reversal persisted with volume-weighted network means (DDI 0.805; both preferences in 0/20 heads), with anatomical node means (DDI 0.719; 0/20; Table S2) and under ROAST (negative salience contrast in 12 of 18 evaluable heads). It reflects the aggregate decision rule, not a general failure of conventional montages.

### 3.3 Separation and individual dose coverage favored different pairs

FA-05 × OM-02 produced both preferences in all 20 heads, with median contrasts of +0.209 V/m in C and +0.195 V/m in S, median DDI 1.561 and median Dice 0.000. The conventional alternative, FA-02 × OM-07, produced both in 19/20 heads (contrasts +0.075 and +0.183 V/m; DDI 0.627; Dice 0.186; Table 2). The difference in separation lay mainly on the control side.

**Table 2.** Regional separability and dose favor different complete pairs.

| **Measure** | **FA-05 × OM-02<br>Nominated pair** | **FA-02 × OM-07<br>Conventional alternative** |
|---|---|---|
| Median regional contrast: C; S (V/m) | +0.209; +0.195 | +0.075; +0.183 |
| Median normalized DDI | 1.561 | 0.627 |
| Both p99 directions, heads | 20/20 | 19/20 |
| Median Dice at half robust peak | 0.000 | 0.186 |
| Network p99: FA; OM (V/m) | 0.284; 0.221 | 0.372; 0.469 |
| Both network targets at floor, heads | 13/20 | 20/20 |
| Anatomical p99: FA; OM (V/m) | 0.301; 0.214 | 0.372; 0.431 |
| Both anatomical targets at floor, heads | 11/20 | 20/20 |
| dACC p99: FA; OM (V/m) | 0.069; 0.105 | 0.252; 0.356 |
| dACC at floor: FA; OM, heads | 0/20; 0/20 | 19/20; 20/20 |
| Suprathreshold volume: FA; OM (cm³) | 12.91; 18.53 | 63.87; 93.15 |
| ROAST DDI; both directions | 1.502; 18/18 | 0.517; 20/20 |

All primary values use SimNIBS at 2 mA and baseline conductivities unless the row states ROAST. C and S are frontal network subsets; anatomical targets are left middle-frontal cortex and strict right anterior insula. The floor is 0.20 V/m. Contrasts are medians of within-head differences. ROAST held ring centers fixed and could not construct OM-02 in two heads.

Dose coverage favored the conventional pair. Both pairs passed the cohort-median rule, but FA-05 and OM-02 reached the network floor together in 13/20 heads and both anatomical targets in 11/20, whereas FA-02 × OM-07 reached both network and both anatomical floors in 20/20 (Figs. 2 and 3).

**Fig. 2.**
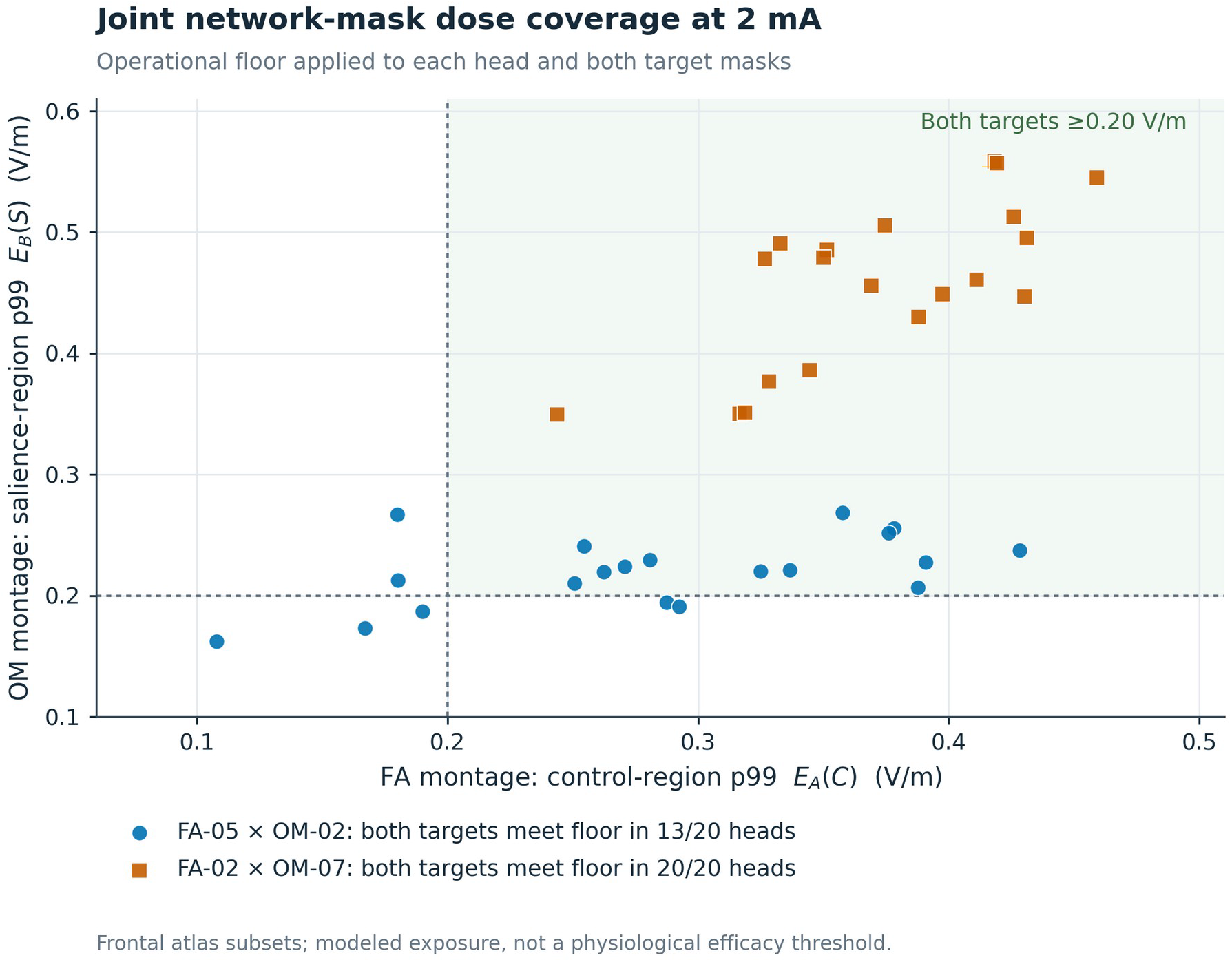
Per-head network dose. Each point gives the two own-target network p99 values for one head; dotted lines mark the 0.20 V/m floor. Joint coverage is 13/20 for FA-05 × OM-02 and 20/20 for FA-02 × OM-07, although both pairs pass the cohort-median dose rule. Regions are frontal atlas subsets; values are modeled exposure at baseline conductivities.

**Fig. 3.**
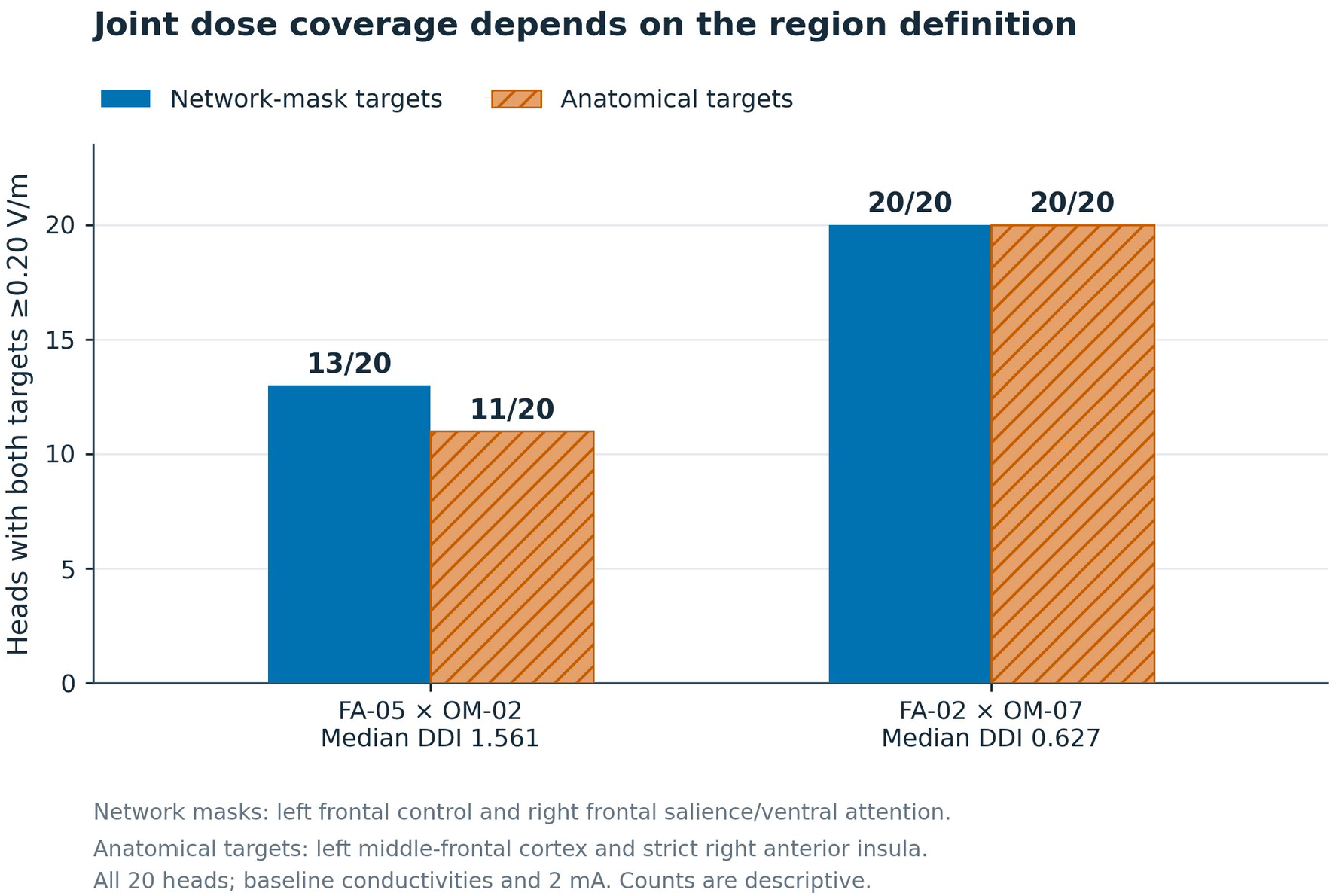
Joint dose coverage by region definition. Both network targets reach 0.20 V/m in 13/20 nominated-pair heads and both anatomical targets in 11/20; the conventional alternative reaches both floors in 20/20 under either definition despite its lower median DDI. Counts are descriptive, at 2 mA and baseline conductivities.

The nomination of FA-05 × OM-02 was therefore a post-results decision to favor separation and restricted suprathreshold exposure, not evidence that it dominates. Pairs were not interchangeable: FA-05 × OM-07 had median DDI 1.132 but both p99 preferences in only 8/20 heads (Table S2).

### 3.4 Restricted exposure did not establish network selectivity

The nominated pair’s separation came with restricted, not selective, exposure. FA-05’s suprathreshold gray-matter volume was 12.91 cm^3^, against 63.87 cm^3^ for FA-02 (OM-02 18.53 versus OM-07 93.15 cm^3^), but its mean field in the anatomical FA target was lower (0.150 versus 0.222 V/m; Table S3).

dACC p99 reached 0.20 V/m in 0/20 heads under either nominated montage and in 19/20 and 20/20 under FA-02 and OM-07; below-floor exposure is not zero exposure. Only 1.5% of the right-hemisphere salience/ventral-attention network exceeded 0.20 V/m under OM-02, versus 79.1% under OM-07, so the frontal-mask contrast cannot be read as selective modulation of the salience network.

### 3.5 Separation was more robust than dose

Both leading pairs met all four recorded rules under ROAST. For FA-05 × OM-02, with ring centers held fixed, median DDI was 1.502 and both preferences held in 18/18 evaluable heads (ROAST could not construct the OM-02 pads in two heads); FA-05’s network p99 was lower than in SimNIBS in every head (median 0.231 versus 0.284 V/m). FA-02 × OM-07 gave DDI 0.517 and both preferences in 20/20 heads. Agreement did not extend to every candidate: two OM-04 sponge pairs crossed the overlap ceiling under ROAST (Supplementary Section S4; Table S4).

Across the 60 archived conductivity stress-test draws, FA-05 × OM-02 kept both preferences on every draw (DDI 5th–95th percentile band 1.33–1.71) but reached both network floors in 19/60, against 34/60 for FA-02 × OM-07 (Table S5). The post hoc subset within the corrected review’s ranges retained 40 draws per pair: both directions held in 40/40 and 39/40, and both network floors in 15/40 and 28/40, respectively (Table S7; Supplementary Section S8). Surface sampling lowered p99 by about 8.5% and put OM-02 below its floor in both evaluated heads (Supplementary Section S5). At normalized 2 mA, exploratory corrected-objective arrays separated more cleanly (DDI about 1.70; Dice 0.000) but reached the network floor in 6/20 and 0/20 heads; normalization did not preserve their original per-electrode constraints (Supplementary Sections S6 and S8). Orbitofrontal configurations for a third practice were assessed separately (Supplementary Section S7).

## 4. Discussion

### 4.1 Aggregate screening rules can pass pairs that fail the experiment’s requirements

The principal result concerns how montage pairs should be screened, not which pair is best. The regional-matching design evaluated here requires each montage to exceed the other in its own target region, but the summed index in our recorded screen did not test that requirement: 13 of the 18 pairs it passed failed the directional conjunction, and FA-02 × OM-02 reversed the intended salience-region preference in every head. The cohort-median dose rule had a parallel limitation; both leading pairs passed it, yet joint network coverage was 13/20 heads for one and 20/20 for the other. Separation and individual coverage also traded off, and separation was more robust than dose to solver and conductivity changes. A screen intended to produce reciprocal regional exposure should therefore prespecify the directional conjunction, report per-head joint dose alongside cohort medians, and judge separation and dose as distinct criteria. These criteria follow from the design, not from meditation, and apply to any pair of montages expected to favor different regions.

Because the directional conjunction was added after the original results were inspected, it demonstrates the issue but cannot retroactively become a prespecified validation of the selected pair. The same restraint applies to the post hoc mapping correction and the reinstatement of OM-07: transparent correction makes the analysis interpretable, but does not erase the role of outcome-informed development.

FA-05 × OM-02 was nominated for its larger control-region contrast, consistent signs and smaller suprathreshold extent; the conventional pair’s superior dose coverage is the strongest argument against that choice. Neither the DDI nor the operational floor supplies a utility function that makes one pair universally preferable; the two pairs embody different priorities.

The same criteria apply to optimized current patterns. Optimization can target weighted cortical patterns directly [21], but our exploratory corrected-objective patterns separated more cleanly than the nominated pair while leaving most heads below the dose floor. Rescaling them to a common 2 mA also exceeded the original 1 mA per-electrode constraint in some heads (Supplementary Section S8). An optimized pattern therefore needs both a per-head dose check and a check that the reported current normalization preserves its delivery constraints.

### 4.2 Regional exposure does not establish network-matched effects

FA and OM share executive, salience and default-mode processes; their proposed difference is one of emphasis [3,5,8]. The mapping from a practice to a preferred montage therefore remains a hypothesis even when the fields differ as intended. Because dACC is itself a salience-network node, a low-dACC-exposure pair and a broader salience pair address different mechanistic questions, and field magnitude does not establish the sign of neuronal modulation, which depends on task state and connectivity [12,13]. Existing stimulation-plus-mindfulness trials show heterogeneous or absent augmentation [9,30,31]; they constrain enthusiasm but do not test the regional matching hypothesis.

### 4.3 Next step: delivery and physiological discrimination

Before any behavioral test, one complete pair’s physical delivery, individual exposure, tolerability, masking and physiological effects should be verified. The F8–Fp2 pads need checks of orientation, separation, contact and saline bridging, and the ring needs documented electrode coordinates and shared-return wiring. Mixed ring and sponge hardware also differs in sensation and expectancy; a hardware main effect cannot by itself produce both directions of a predicted reversal, but a hardware-by-practice interaction could imitate one, so masking and measured practice fidelity are required. Low modeled dose should not trigger post hoc current escalation or single-arm substitution; an alternative pair needs revalidation under a prospectively fixed rule. The full behavioral design is specified elsewhere [32].

### 4.4 Limitations

The sample comprises 20 existing, atlas-mapped head models, not a recruited clinical population, and facial masking in the source models can affect supraorbital geometry [15]. Recorded sex labels from the HCP were used only to balance selection; with ten heads per label, sex-stratified comparisons were not attempted. Tissue properties were fixed at baseline. Conductivity sensitivity used two held-out heads, whereas the surface comparison used two heads from the evaluation set; neither analysis provides a calibrated distribution of participant dose. The conductivity subset is post hoc and retains a subset of the original sampling design, rather than a new Latin hypercube. The placement bar pooled two anchor pairs under the earlier mapping and is not a pair-specific uncertainty estimate. A prospective external cohort and a pair-specific uncertainty analysis remain outstanding.

ROAST used fixed SimNIBS centers for rings, failed two OM-02 constructions and did not re-solve the optimized arrays, and neither pipeline was validated against intracranial measurements in these heads. The p99 and cohort-median summaries, relative-threshold Dice and frontal masks each depend on their definitions. The magnitude summaries do not resolve cortical-normal or tangential field components, cell orientation, or state-dependent response; reciprocal p99 contrasts alone cannot establish reciprocal physiological modulation.

Candidate selection, the directional conjunction, the cortical cap and the corrected optimization were informed by development results and are not preregistered confirmation. The conductivity subset and current-normalization audit added in this revision are also post hoc. The deposited snapshot supports reproduction of the original tabulated analyses [19]; the separate submission reproduction addendum supplies the new calculations and their inputs. A full finite-element rerun and independent physiological validation have not been completed. Independent human methodological critique remains outstanding.

## 5. Conclusions

Summed separation indices and cohort-median dose rules passed montage pairs that failed the intended reciprocal regional-exposure criteria. In the tested set, separation and individual dose coverage favored different pairs, and separation was more robust than dose to the tested solver and conductivity changes. Screens intended to create reciprocal regional exposure should prespecify the directional conjunction and report per-head joint dose. These results describe modeled exposure; they do not establish selective network modulation, practice-specific learning or clinical efficacy.

## Declarations

## CRediT authorship contribution statement

Dylan Bruschi: Conceptualization, Methodology, Software, Validation, Formal analysis, Investigation, Data curation, Writing – original draft, Writing – review & editing, Visualization, Project administration. Ivanna Joseph: Investigation, Writing – review & editing. Adam Schiavi: Conceptualization, Supervision, Writing – review & editing. D.B. configured and executed the current-flow simulations, wrote and maintains the analysis code and configurations, produced the computational results, and is accountable for them; D.B. identified and disclosed the specification defects. I.J. and A.S. reviewed the reported results and their interpretation and did not run the pipeline. No author’s institution independently verified the simulations.

## Ethics and study status

This computational study used existing anatomical head models and recruited no new participants. The models are de-identified, facially masked derivatives published by Berger et al. [15] under CC BY 4.0, built from Human Connectome Project S1200 imaging acquired under the HCP’s own institutional review board approval and participant consent; the sex and age-band labels used for selection came from the HCP’s open-access tier under the WU-Minn HCP Consortium Open Access Data Use Terms, which the corresponding author accepted. No participant is identifiable and no new data were collected. As a secondary analysis of de-identified, publicly released data, the study has not been submitted for an institutional review board determination; none has been sought to date. It reports no intervention, newly collected participant data, or clinical outcomes. The proposed future trial has not been conducted, registered, or funded. Any implementing participant study requires its own appropriate ethics review, independent oversight, prospective registration, and prespecified analysis plan before enrollment.

## Data and Materials Availability

The original analysis is identified by tdcs-modeling commit 0209ed23e017e1e0941f465d3e56140211fef3b6 (tag v1.38-manuscript, 15 October 2026), including progression run 20261005T181232-58a91f22. The primary cohort results, second-solver summaries and original 60-draw conductivity results retain the values reported from that snapshot. It contains the configurations, ROI and montage definitions, analysis code, threshold records, per-head metrics, pairwise results, progression decisions, second-solver outputs, ring electrode coordinates, sensitivity analyses, recorded predictions, and development logs. It does not contain complete meshes, leadfields or raw field arrays. The post hoc conductivity subset and optimized-current normalization audit in Supplementary Section S8 were computed during this revision and are supplied separately in the submission reproduction addendum; they are not part of the original snapshot.

The repository is private at the time of this version. The cited analysis snapshot is deposited publicly on Zenodo ([19]; doi:10.5281/zenodo.23406359). Its MIT license covers code only; head-model-derived data retain their source terms. The original tabulated analyses can be recomputed from the recorded results, and results/versions.json records the archived software environment. The separate submission reproduction addendum contains reproduce_submission_addendum.py, exact copies of the required archived input files, their SHA-256 provenance, and expected outputs for Supplementary Section S8.

Figure plotting code is also supplied with this submission. Editors and reviewers may request read-only access to a history-free copy of the cited commit from the corresponding author at https://github.com/Sunkara-Wellness/tdcs-modeling-review; the copy’s README describes its checks and omissions.

The package supports reproduction of tabulated analyses and figures, not a complete simulation rerun. The latter additionally requires the Berger et al. [15] anatomical models, SimNIBS 4.6, ROAST 4.0 under MATLAB, the solver environments, and substantial computation. Per-head field volumes (about 10 GB) and the solved meshes are not deposited because of their size and because, as derivatives of HCP data, they may be redistributed only under the WU-Minn HCP Consortium Open Access Data Use Terms, which attach to the CC BY 4.0 Berger et al. [18] models as well; they can be regenerated from the deposited configurations with those models, and are available from the corresponding author on request under those terms. Source-model access is governed by the dataset and underlying Human Connectome Project terms. The secondary audit and new figure calculations used recorded outputs, not an independent finite-element rerun. Template validation used the ErnieExtended model [33], supplied under CC BY-NC 4.0; its source terms apply separately from those of the Berger cohort models and analysis code.

## Declaration of generative AI and AI-assisted technologies in the manuscript preparation process

During the preparation of the earlier manuscript, the authors reported using Claude (Anthropic; Claude Opus 5.5 and Claude Fable 5.1) and ChatGPT (OpenAI; ChatGPT 6 Astra) to edit, restructure and revise text and assist literature retrieval. During the present submission revision, ChatGPT (OpenAI; GPT-6 Astra Pro) assisted literature verification, scientific editing, the numerical audit, reproducible supplementary calculations and figure plotting. Responsibility for the content, analysis and interpretation remains with the named authors. No AI system is an author. Code assistance is described in Section 2.7 and Supplementary Sections S1 and S8.

## Acknowledgments

The authors thank collaborators and advisors at Sunkara Wellness Technologies for discussion of the network model and experimental design, and acknowledge Berger et al. [15] for the anatomical models. Data were provided in part by the Human Connectome Project, WU-Minn Consortium (Principal Investigators: David Van Essen and Kamil Ugurbil; 1U54MH091657) funded by the 16 NIH Institutes and Centers that support the NIH Blueprint for Neuroscience Research; and by the McDonnell Center for Systems Neuroscience at Washington University.

